# A Single-cell Transcriptomic Atlas of the Developing Chicken Limb

**DOI:** 10.1101/598227

**Authors:** Christian Feregrino, Fabio Sacher, Oren Parnas, Patrick Tschopp

## Abstract

**Background:** Through precise implementation of distinct cell type specification programs, differentially regulated in both space and time, complex patterns emerge during organogenesis. Thanks to its easy experimental accessibility, the developing chicken limb has long served as a paradigm to study vertebrate pattern formation. Through decades’ worth of research, we now have a firm grasp on the molecular mechanisms driving limb formation at the tissue-level. However, to elucidate the dynamic interplay between transcriptional cell type specification programs and pattern formation at its relevant cellular scale, we lack appropriately resolved molecular data at the genome-wide level. Here, making use of droplet-based single-cell RNA-sequencing, we catalogue the developmental emergence of distinct tissue types and their transcriptome dynamics in the distal chicken limb, the so-called autopod, at cellular resolution.

**Results:** Using single-cell RNA-sequencing technology, we sequenced a total of 17,628 cells coming from three key developmental stages of chicken autopod patterning. Overall, we identified 23 cell populations with distinct transcriptional profiles. Amongst them were small, albeit essential populations like the apical ectodermal ridge, demonstrating the ability to detect even rare cell types. Moreover, we uncovered the existence of molecularly distinct sub-populations within previously defined compartments of the developing limb, some of which have important signaling functions during autopod pattern formation. Finally, we inferred gene co-expression modules that coincide with distinct tissue types across developmental time, and used them to track patterning-relevant cell populations of the forming digits.

**Conclusions:** We provide a comprehensive functional genomics resource to study the molecular effectors of chicken limb patterning at cellular resolution. Our single-cell transcriptomic atlas captures all major cell populations of the developing autopod, and highlights the transcriptional complexity in many of its components. Finally, integrating our data-set with other single-cell transcriptomics resources will enable researchers to assess molecular similarities in orthologous cell types across the major tetrapod clades, and provide an extensive candidate gene list to functionally test cell-type-specific drivers of limb morphological diversification.

## Background

Embryonic pattern formation relies on the tight coordination of numerous developmental processes, across multiple scales of complexity. From seemingly homogenous progenitor populations, different cell types get specified and arranged in intricate patterns, to give rise to functional tissues and organs. As progenitors mostly share a common genome, this phenotypic specialization relies on the precise execution of distinct gene regulatory networks, to enable cell type specification and ensuing pattern formation [1–3]. Slight deviations in these processes contribute to morphological variations within natural populations. More profound aberrations, however, can cause malformations and ultimately result in death of the embryo. To buffer such fragile balance, many cell type specification and pattering processes rely on complex feedback mechanisms, through tightly interconnected molecular loops between spatially distinct signaling centers [4–6] Hence, integration of multiple signaling pathways across space and time define a molecular coordinate grid to instruct organogenesis at the tissue level. Ultimately, however, these multifaceted signaling inputs have to be incorporated at the cellular level, *via* cell type-specifying gene regulatory networks, as progenitor cells undergo spatially and temporally defined cell fate decisions to contribute to proper pattern formation.

Tetrapod limb development has long served as a model to study the genetic and molecular underpinnings of vertebrate pattern formation. Due to its non-essentiality for embryo survival, many fetuses carrying mutations that affect limb development make it to full term. Accordingly, human geneticists have been able to accumulate an impressive catalogue of candidate genes for limb patterning [7–9]. Combined with the easy accessibility of the limb in chicken embryos, and molecular genetic tools in the mouse, decades of experimental work have resulted in an in-depth understanding of many of the molecular mechanisms driving limb formation at the tissue scale [5]. Moreover, given the profound morphological diversifications the basic limb structure has experienced in numerous tetrapod clades, limb development has long attracted the interests of comparative developmental biologists using ‘EvoDevo’ approaches [10]. This holds especially true for the most distal portion of the limb, the autopod, i.e. hands and feet. There, species-specific adaptations to distinct modes of locomotion have resulted in a diverse array of digit number formulas and individualized digit patterns [11–14].

Early in development, proliferation of a lateral plate mesoderm (LPM)-derived mesenchymal progenitor population drives overall limb bud outgrowth. Signaling crosstalk with a specialized structure of the distal overlaying ectoderm, the apical ectodermal ridge (AER), controls these dynamics. Concurrently, the major embryonic axes of the limb are defined by the coordinated action of multiple signaling centers [reviewed in 5]. As development progresses, LPM-derived progenitors start to differentiate into skeletal and other connective tissue types [15–17], while muscles cells originating from the somites migrate into the limb bud to complement formation of the musculoskeletal apparatus [18, 19]. For autopod pattern formation, digit numbers and identities are first defined by posteriorly restricted sonic hedgehog (SHH) activity, and altered by modulations therein [10, 14, 20, reviwed in 21]. Digit elongation then relies on a specialized distal progenitor population, which supports outgrowth of individual digit bones, the phalanges [22, 23]. Digit-specific phalanx-formulas, and their stereotypic connection patterns *via* synovial joints, are established by signals emanating from the posterior interdigit mesenchyme [24, 25].

In this study, capitalizing on the power of droplet-based single-cell RNA-sequencing, we resolve the underlying transcriptional dynamics of autopod tissue formation and pattern emergence at single-cell resolution, across three stages of chicken hindlimb development. In total, we present transcriptomic data for 17,628 cells, allowing us to identify all major tissue types of the developing limb, as well as a substantial amount of molecular heterogeneity therein. Through weighted correlation network analysis, we define distinct gene co-expression modules that track corresponding tissue types across developmental time. Finally, we focus on the molecular make-up of cell populations involved in digit pattern formation and, hence, putative drivers of morphological diversification in the autopod.

Collectively, we present a comprehensive genomics resource that for the first time reveals the transcriptome dynamics of the developing chicken foot at cellular level. Our study identifies novel and known marker genes in co-expression modules of patterning-relevant cell populations, thereby providing an extensive catalogue of candidate genes for functional follow-up studies, to elucidate the molecular mechanisms of autopod pattern formation and diversification.

## Results

### Singe-cell sampling of the developing distal chicken limb

To follow the appearance of patterning-relevant cell populations and their associated transcriptome dynamics, we sampled three developmental stages of the embryonic chicken foot: stage Hamburger-Hamilton 25 (HH25, ∼4.5 days of development), stage HH29 (∼6 days of development) and stage HH31 (∼7 days of development). This time window spans key morphogenetic events that drive species-specific patterns in the developing autopod, particularly for the skeletal apparatus and its associated tissues. Namely, stage HH25 is dominated by overall autopod outgrowth and delineation of the main embryonic axes, at HH29 digit-specific patterns differentiate, and at HH31 digit elongation is phasing out. We designed our tissue sampling strategies accordingly. At HH25, we captured the entire distal part of the growing limb (Fig. 1a), at HH29 we dissected two digits with distinct skeletal formulas, digit 3 and 4, as well as their adjacent interdigit mesenchyme (Fig. 1b), and at HH31 we focused on the tip of digit 4 with its growth-relevant progenitor population (Fig. 1c). We dissociated the micro-dissected tissue pieces using enzymatic digest combined with mechanical shearing and prepared single-cell suspensions for droplet-based high-throughput single-cell RNA-sequencing (*10X Genomics* and *Drop-Seq* [26, 27]). Using the corresponding bioinformatics pipelines, the resulting Next-Generation Sequencing libraries were mapped to the chicken genome, de-multiplexed according to their cellular barcodes and quantified to generate gene/cell read count tables. In total, we sampled over 17,000 cells and obtained single-cell transcriptomic profiles for 5,982 (HH25), 6,823 (HH29) and 4,823 (HH31) individual cells, respectively (Additional file 1: Fig. S1a). Quality-based exclusion of single-cell transcriptomes was implemented based on mean library size, percentage of mitochondrial reads and number of genes detected per cell. Additionally, data normalization as well as batch and cell cycle corrections were performed (for details, please refer to the *Methods* section). On average, we detected 2,879 unique molecular identifiers (UMIs) and 1,081 genes per cell (Additional file 1: Fig. S1b,c).

**Fig. 1.**
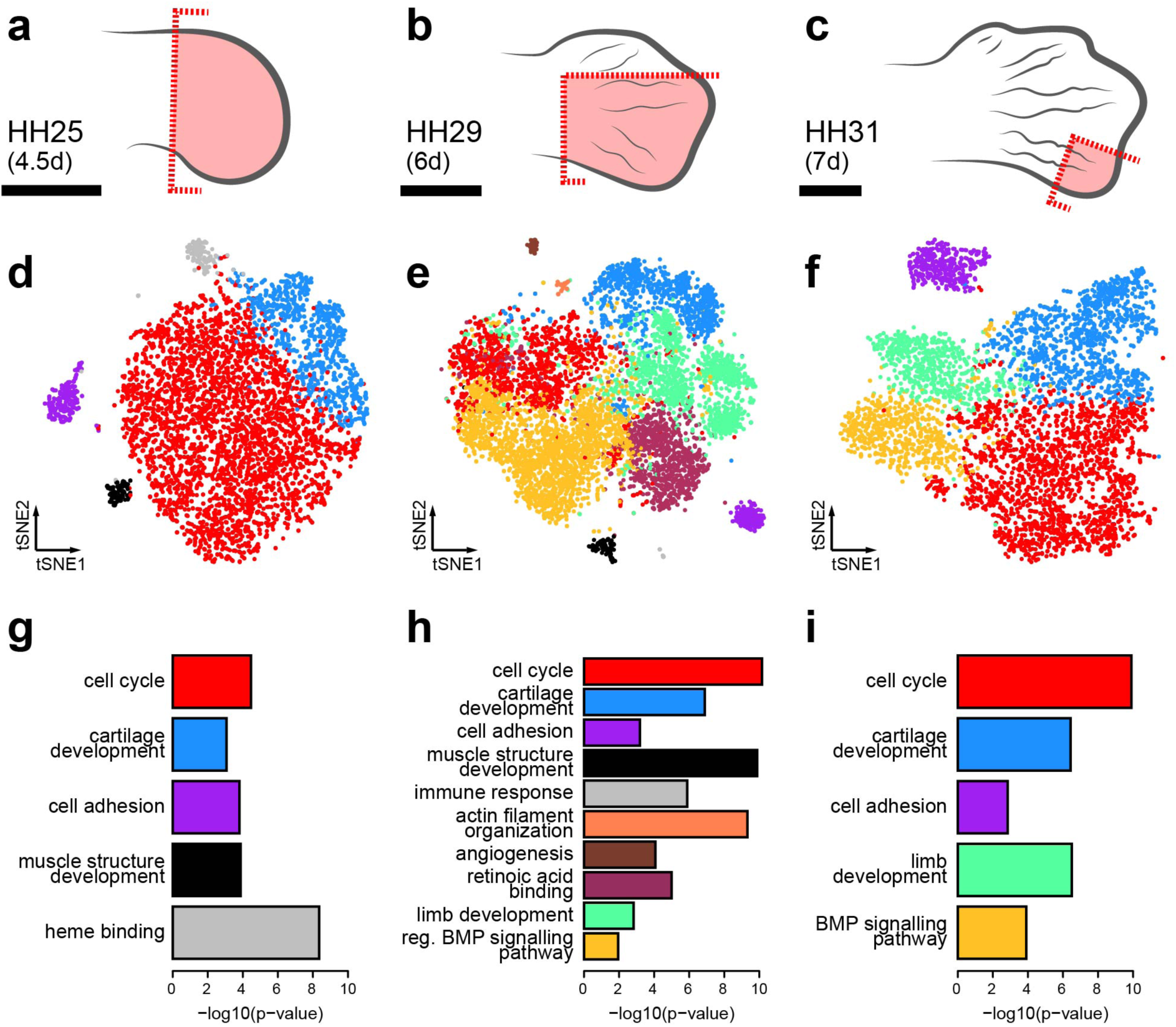
Sampling strategy and tissue composition of the developing chicken autopod. **(a-c)** Dissection schemes, highlighted in red, for sampling the different stages of hindlimb development (scale bar ∼1mm). **(d-f)** tSNE representation of the three datasets, representing 5,982 (HH25), 6,823 (HH29) and 4,823 (HH31) according to their transcriptome similarities. Cellular color codes reflect unsupervised graph-based clustering results. Comparable cell populations identified in multiple samples are visualized using the same color. **(g-i)** Select overrepresented GO-terms, from analysis of the overexpressed genes, for each cluster at stages **(g)** HH25, **(h)** HH29 and **(i)** HH31.

### Autopod tissue composition at cellular resolution

Using unsupervised graph-based clustering, we identified 5, 10 and 5 clusters at stages HH25, HH29 and HH31, respectively. Projecting these clusters onto stage-specific tSNE (t-Distributed Stochastic Neighbor Embedding [28]), plots of our cellular transcriptomes revealed the presence of a dominant bulk of cells, with varying degrees of sub-structure, as well as distinct outlier groups (Fig. 1 d-f). Based on the expression of known marker genes and gene ontology (GO)-term enrichment analyses, we were able to attribute these broadly defined cell populations to distinct tissue types (Fig. 1g-f, Additional file 1: Fig. S1a and Fig. S2a-c). At stage HH25, they comprise a largely undifferentiated and proliferating mesenchymal population (red), early skeletal progenitors (blue), muscle cells invading the limb (black), as well as skin (purple) and blood cells (grey) (Fig. 1d,g). We recovered cell populations corresponding to those same five tissue types in our HH29 sample, with the exception that the “blood cluster” was now dominated by white blood cells and not erythrocytes. Additionally, we identified cell populations matching the interdigit mesenchyme (green), non-skeletal connective tissue (nsCT, maroon), cells enriched for markers of the very distal margin of the autopod mesoderm (“distal mesenchyme”, yellow), as well as endothelial (brown) and smooth muscle (orange) cells of the forming blood vessels (Fig. 1e,h). At stage HH31, we again find a largely undifferentiated mesenchymal population, the interdigit and distal margin mesenchyme, skeletal and skin cells (Fig. 1f,i). As expected according to our sampling strategy, for spatial and/or temporal context, we did not find all cell populations in every dataset. For example, while sample HH25 is biggest in relative size to the autopod, it is the earliest stage and thus predictably displayed the lowest cellular complexity. We observed the opposite trend in HH31, where the relative size is smallest but development more advanced. Our most complex dataset, in terms of cell number and tissue types identified, is from stage HH29. Collectively, using broad graph-based clustering and molecular profiling on our single-cell transcriptomics data, we catalogued the tissue composition of the developing autopod with cellular resolution, across three developmental stages.

### Fine-scale clustering and marker gene expression across developmental time

Although all expected major tissue types were recovered in our primary analyses, smaller cell populations, some well known to be essential for limb outgrowth and patterning, remained elusive. Hence, given our sampling depth, we next examined our data for additional sub-structure. Indeed, upon closer inspection using finer-tuned clustering parameters, we did find additional sub-populations with distinct transcriptional signatures (Fig. 2a-c, Additional file 1: Fig. S1a). Based on differential expression analyses, we identified marker genes for each of these sub-populations (Additional files 2-4). Certain sub-population/marker gene-combinations appeared to be conserved in all three samples, thereby allowing us to assign cellular equivalencies across developmental time (Fig. 2d-f). A subset of marker genes only showed loosely restricted expression patterns, likely a reflection of the largely undifferentiated state of the corresponding sub-population. For example, *PRRX1*, a well-established marker of the limb mesenchyme [16, 29, 30], and *PCNA*, active during DNA replication in proliferating cells [31], showed varying levels of expression beyond the proliferating mesenchyme sub-clusters. Such transcriptional ambiguities, however, seemed progressively lost, as mesenchymal progenitors committed to the different skeletal and non-skeletal lineages that define the emerging autopod patterns (Fig. 2d-f). As expected, cell sub-populations residing outside the LPM-lineage showed more pronounced transcriptome individualizations. For example, at HH25 the ectodermal ‘skin’ population got split into two distinct sub-clusters, one representing the bulk amount of the embryonic skin covering the autopod (sub-cluster 8), and the other corresponding to the apical ectodermal ridge (sub-cluster 7). Expression of its canonical marker *FGF8* and other highly enriched genes clearly established AER identity, demonstrating that even small cell populations can be successfully captured (Fig. 2d).

**Fig. 2.**
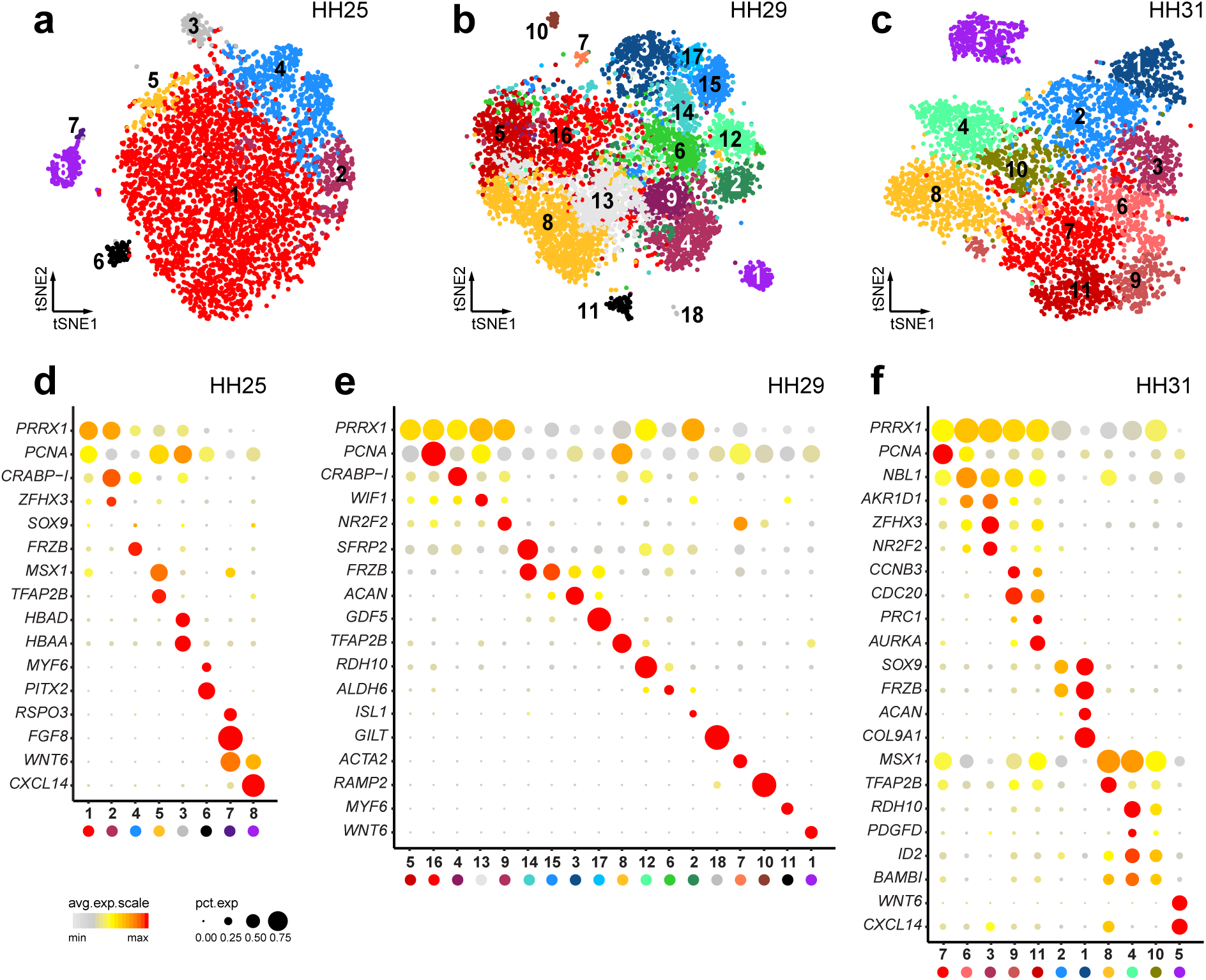
Cell population sub-structure and marker gene expression. **(a-c)** tSNE plots of the three datasets. Colors now represent fine-tuned unsupervised graph-based clustering, with similar colorations relating to the results of the first clustering step. Comparable cell populations identified in multiple samples are visualized using the same color. For reference, sub-cluster numbers are added. **(d-f)** Dot plots of sub-cluster marker gene expression. Averaged expression level (heatmap) and percentage of cells showing >0 expression (dot size) is visualized across all samples, for all identified sub-clusters. Same color-coding for sub-clusters identification is used as in **(a-c)**.

### Gene co-expression modules and corresponding tissue types

To gain further insights into the regulatory programs that maintain these transcriptional signatures, and explore their potential biological significance, we tested for the occurrence of transcriptome-wide gene co-expression patterns using weighted correlation network analysis (WGCNA) [32]. This approach consists of an unsupervised clustering of genes based on their expression pattern across all cells, irrespective of the assigned cell or tissue type. In order to comprehensively screen for relevant gene co-expression modules, we conducted the analysis in our transcriptionally most complex sample at stage HH29. Starting with genes that showed high levels and variation of expression, we calculated an adjacency matrix and its topological overlap to construct a hierarchical tree. The resulting tree was cut to obtain a first set of gene co-expression modules. We then computed the first principal component of each module, to define so-called ‘module eigengenes’. For each individual gene, correlation to the respective eigengenes was used to assess module membership. Genes not significantly correlated with any eigengene were discarded, after which the entire process was repeated iteratively with a reduced gene set. Eventually, we identified a total of 836 genes grouped in 16 distinct gene co-expression modules, each designated by a color (Fig. 3a). Final module sizes ranged from 15 to 215 genes (Additional file 5).

**Fig. 3.**
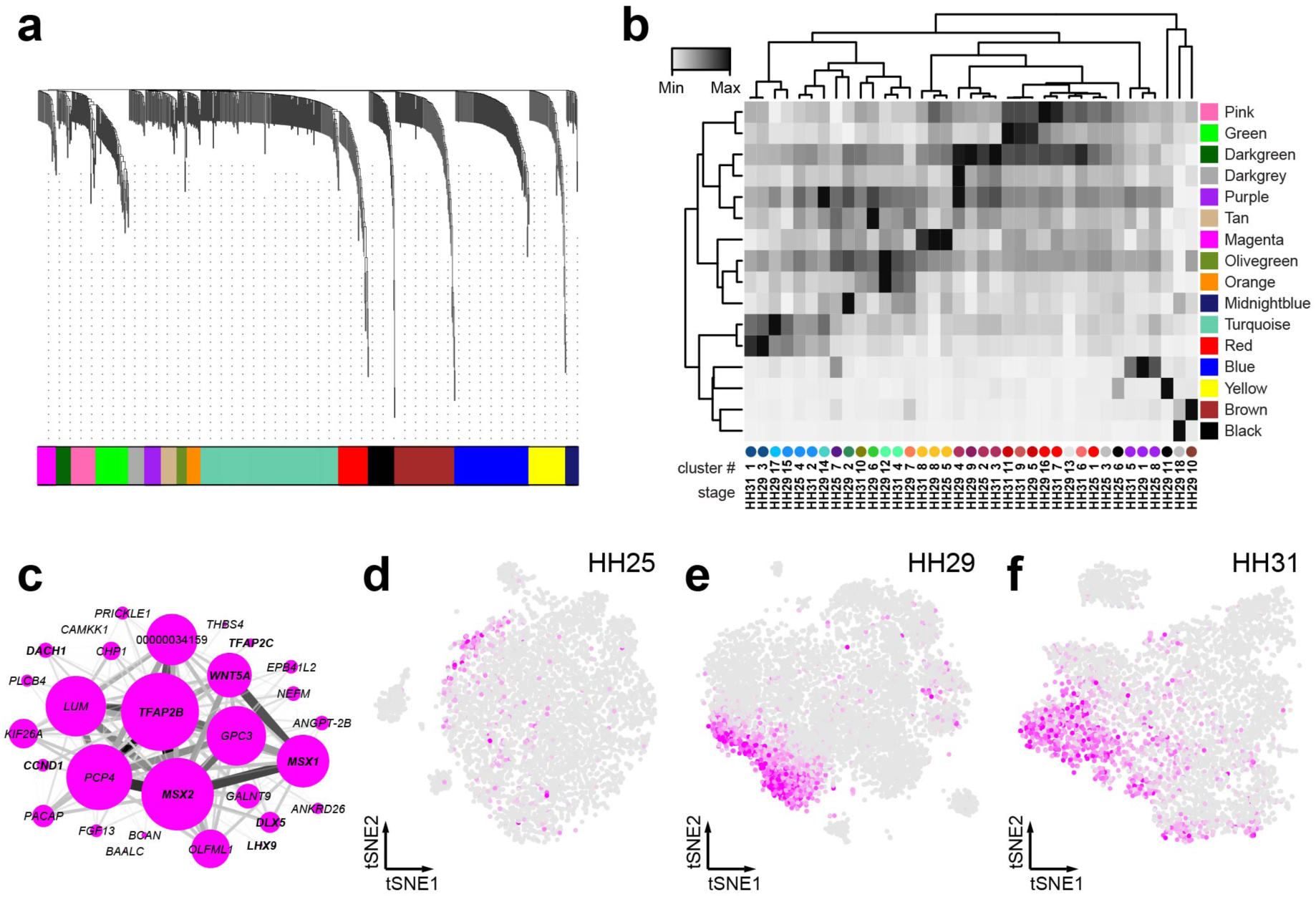
Weighted correlation network analysis and gene co-expression modules. **(a)** WGCNA gene hierarchical clustering dendrogram and modules of co-expression. A total of 16 distinct co-expression modules are identified, visualized by colored bars at the bottom of the dendrogram (color scheme unrelated to previous cell clustering). **(b)** Heatmap of mean expression values per co-expression module, calculated across distinct cell sub-clusters and developmental stages. Ordering based on hierarchical clustering of averaged co-expression module activities and sub-clusters. Sub-clusters identification at bottom (number and color code) corresponds to Fig. 2a-c. **(c)** *Cytoscape* visualization of co-expression module Magenta. Node size is proportional to module membership of each gene, edge thickness represents correlation of pair-wise gene co-expression. **(d-f)** Heatmap representing the averaged cellular activity of the Magenta module, plotted on tSNE representations of the different samples. Color intensity is proportional to the mean expression of the module in each cell.

On a cell-by-cell basis, we calculated the average expression for each of the co-expression modules and visualized their distribution on our stage HH29 tSNE plot (Additional file 1: Fig. S3). Compared to our initial clustering of sample HH29, we found co-expression modules specifically enriched in the following cell populations: blood cells (module Black), skin (Blue), blood vessel endothelium (Brown), nsCT (Darkgrey), distal mesenchyme (Magenta), chondrocytes (Red and Turquoise) and muscle (Yellow). Interestingly, GO-terms associated with more broadly distributed modules enabled us to attribute the sub-clustering structure of certain tissues to particular biological processes. For example, HH29 mesenchyme sub-cluster 5 showed higher activity for module Green, associated with GO-terms connected to mitosis, whereas sub-cluster 16 was enriched for module Pink, linked to G2/M-transition-related genes (Additional file 1: Fig. Fig. S3). Hence, we reasoned that distinct cell-cycle states underlie the subdivision of the proliferating mesenchyme cluster. Likewise, HH29 interdigit sub-clusters 2, 6 and 12 were closely matched by the activities of modules Tan, Olivegreen, Orange and Midnightblue (see below, Fig. 4a-h).

**Fig. 4.**
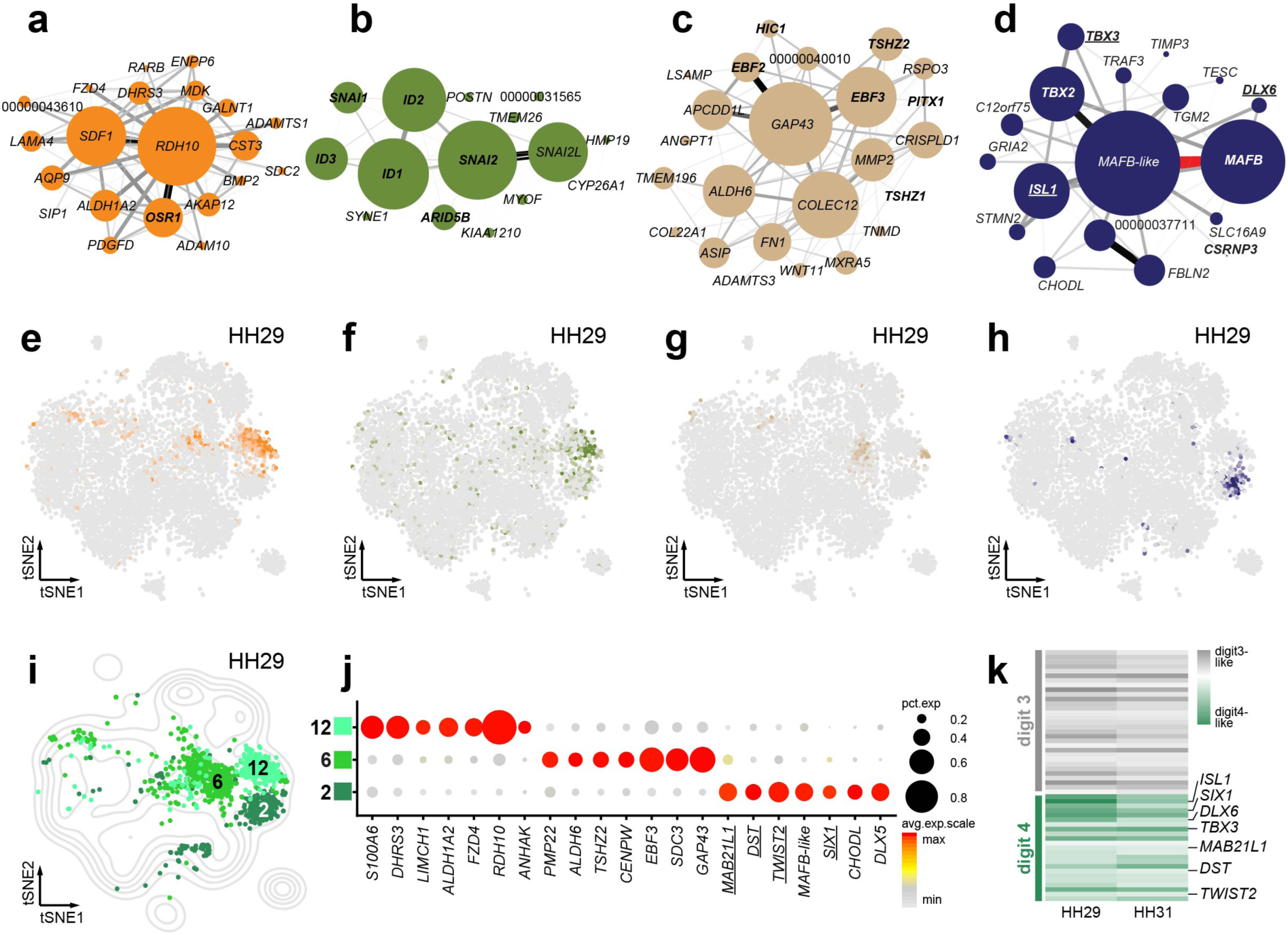
Molecular and spatial heterogeneity in the interdigit mesenchyme. **(a-g)** Interdigit-associated co-expression modules **(a)** Orange, **(b)** Olivegreen, **(c)** Tan, and **(d)** Midnightblue. Node size represents gene module membership, edge thickness gene pair-wise correlation. Gene names in bold are classified as transcription factors, uncharacterized genes show only Ensembl numbers following the “ENSGALG” gene code. **(e-h)** Heatmaps of averaged activity levels of the corresponding modules, visualized on top of a tSNE plot for sample HH29. **(i)** Contour density plot of the tSNE projection for sample HH29, to delineate overall cell distribution. Partial tSNE plot on top, to visualize only cells belonging to interdigit-like sub-clusters (Color-coding and numbering according to Fig. 2b). **(j)** Expression dot plot of differentially expressed genes between the three interdigit sub-clusters at stage HH29. **(k)** Heatmap visualization of “digit3-like” and “digit4-like” gene sets at stages HH29 and HH31, based on differential expression analysis of digit-specific bulk RNA-seq data by Wang et al., 2011. Underlined gene names in **(d,j)** denote membership to the “digit IV-like” gene set.

To follow the developmental dynamics of the identified modules, we calculated their averaged activities across all the three sampled time points, and visualized similarities across time and tissue types using unsupervised hierarchical clustering (Fig. 3b). Indeed, despite differences in embryonic stages and experimental platforms, we were able to confirm corresponding cell and tissue types between our samples. For example, what we refer to as the “distal mesenchyme” is a population of cells characterized by high activity of the co-expression module Magenta at all time points (Fig. 3c-f). Comparisons to published expression patterns for *TFAP2B, WNT5A*, *MSX1* and *MSX2* confirmed its distal location and, based on those genes’ functions, suggested a role for this cell population in controlling distal autopod outgrowth. Using WGCNA thus enabled us to define equivalent cell populations across developmental time, and helped attribute biological functions at the sub-cluster level.

### Transcriptionally and spatially distinct sub-populations in the interdigit mesenchyme

As expected by developmental stage, interdigit populations were only recovered in samples HH29 and HH31. In total, we identified four associated co-expression modules (Fig. 4a-d). High Orange and Olivegreen module activities were coinciding with the same interdigit sub-population (Fig. 4e,f), which was recognizable in both HH29 and HH31 samples and marked by *RDH10* expression (Fig. 2e,f). Noticeably, all genes with high membership in module Olivegreen were transcription factors (TFs), while module Orange was enriched for enzymatic activities (Fig. 4a,b). Both, however, scored high for GO-terms related to retinoic acid signaling, an important mediator of interdigit cell death [33]. Module Tan was enriched for skeletogenic and morphogenetic GO-terms, suggesting it might mediate some of the patterning information contained in the interdigit mesenchyme to the adjacently forming digits (Fig. 4c,g). Lastly, module Midnightblue showed multiple TFs and its activity was restricted to HH29 sub-cluster 2 (Fig. 4d,h).

Since relevant patterning information is contained in the interdigit, posteriorly adjacent to each forming digit, we next wondered whether some of the sub-clustering structure corresponded to spatially distinct interdigit populations along the anterior-posterior axis of the autopod. At HH29, we detected three interdigit sub-clusters (Fig. 4i). Using differential expression analyses, we defined marker genes that distinguish the three sub-clusters from each other (Fig. 4j). To assign putative spatial information to our single-cell interdigit transcriptomes, we reanalyzed a bulk RNA-seq dataset covering stages HH29 and HH31 of the developing chicken hindlimb autopod [34]. This dataset is based on dissections of individual digits, together with their posteriorly associated interdigit mesenchyme, and thus provided an opportunity to identify spatially resolved marker genes. We contrasted their transcriptomic data of digit/interdigit III against digit/interdigit IV and found a total of 54 genes to be significantly differentially expressed at both developmental time points (Fig. 4k). Comparing the digit/interdigit IV-specific subset of these genes to our differential expression analysis of sub-cluster 2, and its affiliated module Midnightblue, we found an overlap of seven up-regulated genes (Fig. 4d,j, underlined). In contrast, we couldn’t find any other digit/interdigit IV gene in the rest of the interdigit sub-cluster signatures or co-expression modules. We therefore concluded that HH29 sub-cluster 2 consisted of cells of the interdigit mesenchyme posterior to digit 4.

### Developing digits and their associated tissues

Of the cell populations directly contributing to the making of digits, a cluster reminiscent of the non-skeletal connective tissue, the nsCT, appeared in all of the samples. In our WGCNA analyses, we identified three modules, Darkgrey, Purple, and Darkgreen, which mapped to the nsCT sub-clusters (Fig. 5a-f). The Darkgrey module was most restricted, in both time and cell numbers, and its activity pattern closely matched the HH29 sub-cluster 4 (Fig. 5d). Cellular retinoic acid binding protein I *CRABP-I*, Aquaporin *AQP1*, *DKK2* and *GLT8D2* were the genes most strongly associated with this module. Modules Purple and Darkgreen showed more widespread activities (Fig. 5e,f), and centered on *COL1A2*, *DCN*, *KCNJ2*, *SALL1*, and *AKR1D1*, *PRRX1*, *TCF12, ZFHX3*. Comparing our differential expression analyses between the respective cell populations, only six genes appeared significantly enriched across all stages (Fig. 5g), five of which also appeared in our nsCT modules. Using *in situ* hybridization for the top-three of these genes, both differential expression- and module membership-wise, allowed us to attribute module activities to discrete nsCT domains along the developing skeletal elements. *CRABP-I* showed highest expression near and around the forming epiphysis, where synovial joints and ligament attachment sites develop (Fig. 5h). *COL1A2*- and *ZFHX3*-positive populations showed a graded distribution along the periskeletal tissue layer, predominantly marking the prospective periosteum and perichondrium domains, respectively (Fig. 5i,j).

**Fig. 5.**
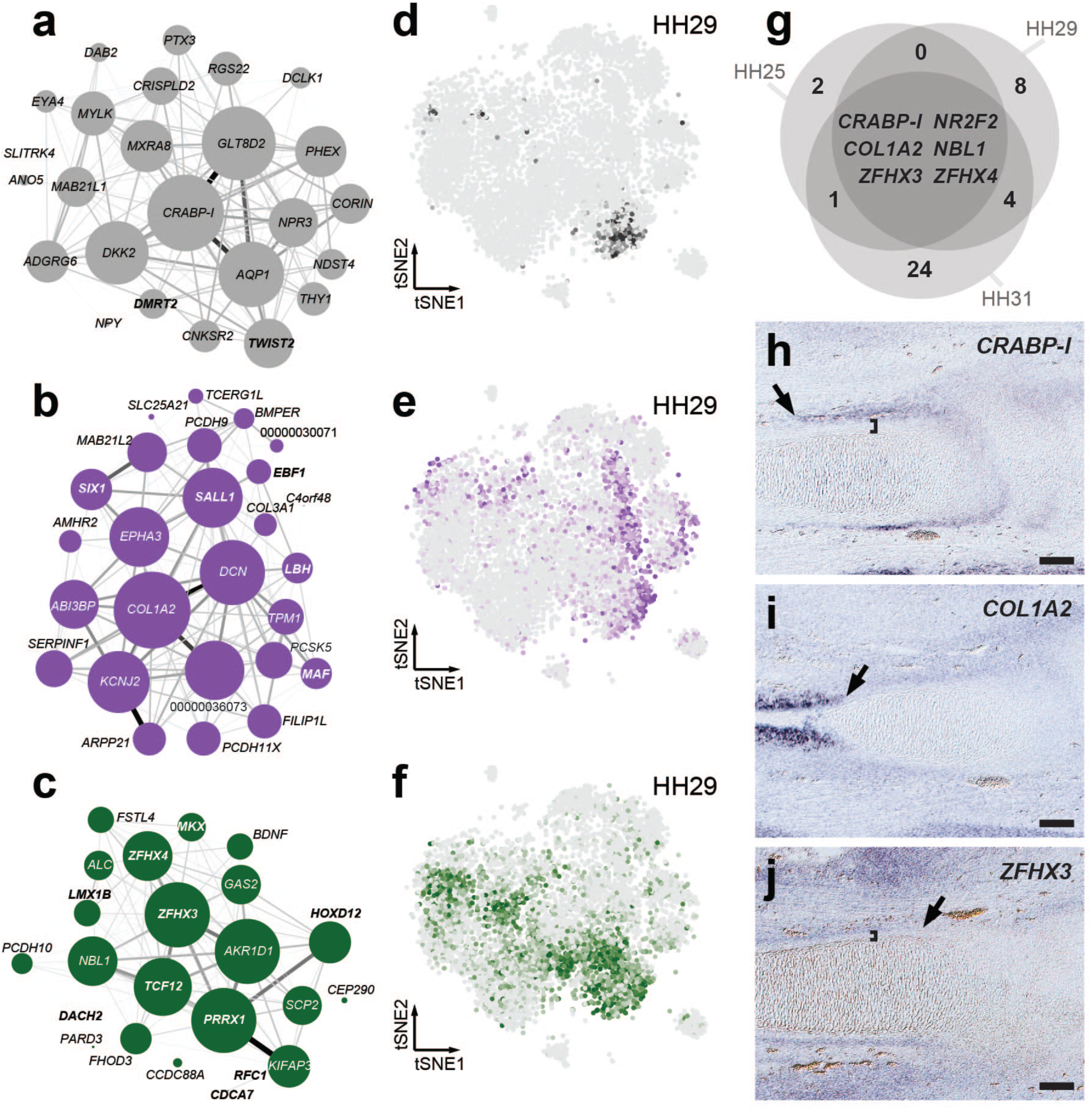
Transcriptional modules in the non-skeletal connective tissue (nsCT). **(a-c)** Gene co-expression modules **(a)** Darkgray, **(b)** Purple and **(c)** Darkgreen enriched for peri-skeletal genes. Gene names in bold are classified as transcription factors, uncharacterized genes show only Ensembl numbers following the “ENSGALG” gene code. **(d-f)** Corresponding averaged module activities visualized as heatmaps on stage HH29 tSNE plots. **(d)** Venn diagram of shared overexpressed genes in the nsCT populations of the three samples. **(h-i)** Section *in situ* hybridization on stage HH31 chicken hindlimbs for three shared nsCT marker genes, *CRABP-I*, *COL1A2* and *ZFHX3*. Arrows denote extent of expression along the long bone axis, while brackets indicate separation from the forming skeletal element (scale bar=100mm).

Finally, we identified skeletal progenitor populations at all three time points (Fig. 6a-c). According to the developmental stages we sampled, only cartilage-producing skeletal cells were recovered. In all three samples, we found a cell population resembling early chondrocytes (sub-clusters HH25-4, HH29-15 and HH31-2). At stages HH29 and HH31, a seemingly more mature chondrocyte type emerged (HH29-3, HH31-1), and an additional cartilaginous cluster was evident in the HH29 sample (HH29-17). Concomitantly, we identified two co-expression modules associated with these cell populations, Turquoise and Red (Fig 6d,e). Turquoise is centered on *CD24*, *CHGB* and *SULF1*, whereas module Red displays a core of collagens *COL9A1* and *COL9A3*, *MATN4*, *C9H2ORF82* (also known as *SNORC* in mammals), and *ACAN*. Based on additional marker genes and GO-term enrichment analyses, we inferred the Turquoise module to be related to early chondrocyte proliferation and growth, whereas the Red module reflected chondrocyte maturation and extracellular matrix deposition (Fig. 6f). Interestingly, compared to module Turquoise, the activity of module Red was generally more restricted and specifically excluded from sub-cluster HH29-17 (Fig. 6g,h). Upon closer inspection, we identified high expression of several known synovial joint markers genes in this population, thus identifying it as the forming interphalangeal joints (Fig. 6i, Additional file 3).

**Fig. 6.**
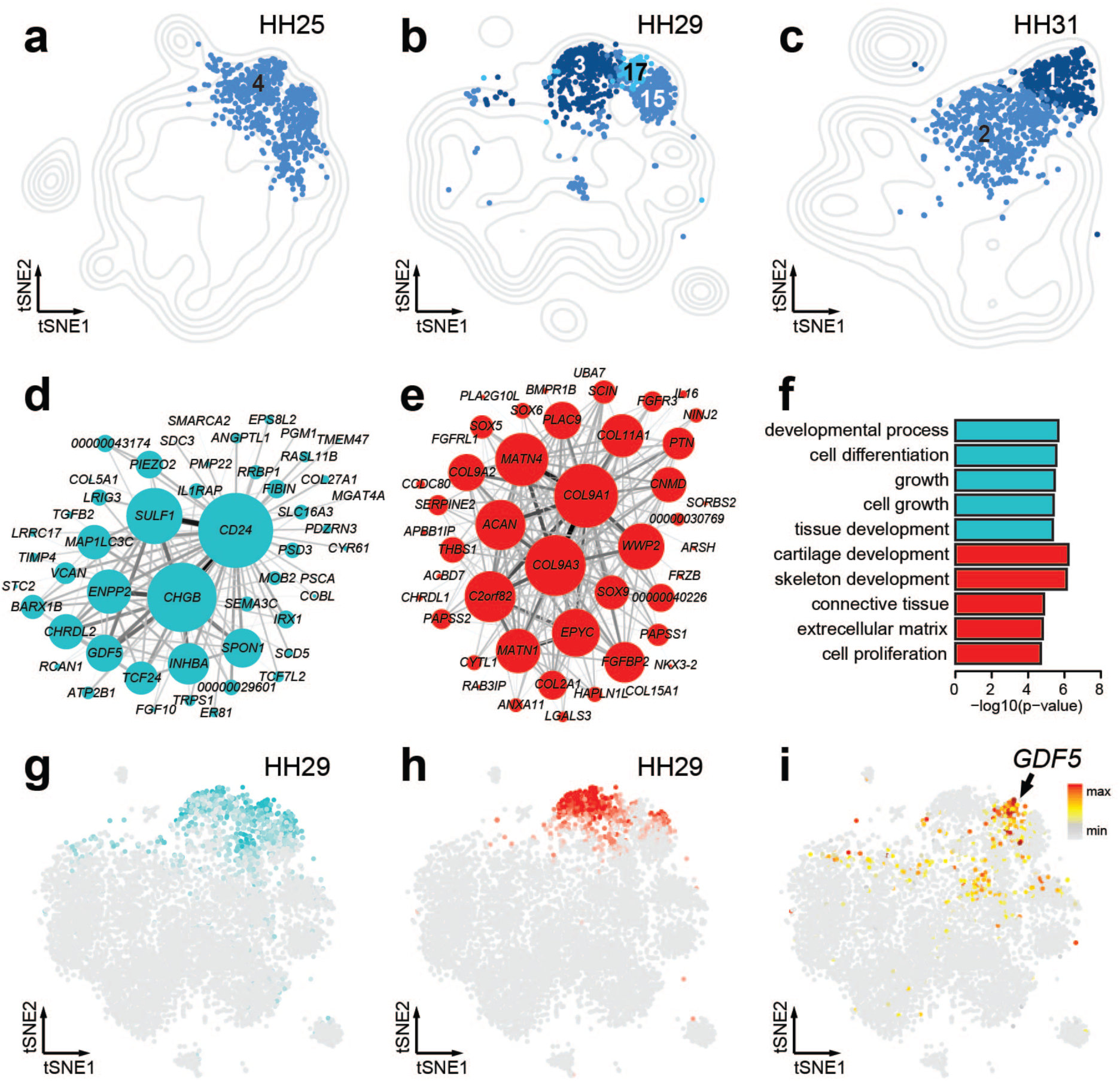
Transcriptional modules and sub-populations in skeletogenic cells. **(a-c)** Contour density plot of tSNE projection for each sample. Partial tSNE plot on top, to visualize only cells belonging to skeletogenic sub-clusters (Color-coding and numbering according to Fig. 2b). Same color / shade across samples indicates comparable cell populations. **(d-e)** Gene co-expression modules **(d)** Turquoise and **(e)** Red. Representation of the Turquoise module only shows the 50 genes with the top membership, of a total of 215. **(f)** Top 5 GO-terms, from analysis of the genes member of modules Turquoise and Red. **(g-h)** Averaged module activities visualized as heatmaps on stage HH29 tSNE plots corresponding to the modules Turquoise and Red. **(i)** Expression heatmap of *GDF5* visualized on stage HH29 tSNE.

Hence, through a combination of differential gene expression and GO-term enrichment analyses, as well as gene co-expression modules, we identified spatially and/or temporally distinct sub-populations and transcriptome dynamics in the skeletal and peri-skeletal tissues of the forming digits.

## Discussion

### Singe-cell tissue decomposition of the developing chicken autopod

Here, using single-cell RNA-sequencing, we present a transcriptomic atlas of the developing chicken limb at cellular resolution. Focusing on the distal and morphologically diverse portion of the limb, the autopod, we sampled over 17,000 single-cell transcriptomes with an average of over 1,000 genes detected in each cell. Within our atlas, we identify all major tissue types that constitute and pattern the embryonic appendage across three developmental time points. Additionally, taking advantage of our cellular and transcriptomic sampling depth, we manage to isolate even minute cell populations like the AER and identify novel marker genes in it. We also distinguish transcriptionally discrete sub-populations within known major tissue types, reflecting distinct spatial locations or cellular states. As such, it demonstrates the power of scRNA-seq to molecularly disentangle cell populations of the developing limb that occur in close spatial or ‘lineage’ proximity. Historically, such populations have proven notoriously difficult to separate and characterize transcriptionally, using either manual tissue dissection or reporter-gene based cell lineage isolation. To what extent all of our tissue sub-clusters indeed correspond to distinct lineage separations [35], or rather represent the extremes of a molecular continuum that follows the inherently stochastic nature of transcription [36, 37], remains to be addressed in future studies. Regardless, however, our results provide a toolbox of candidate genes to tackle this question in a molecularly comprehensive manner. Furthermore, our data enables a characterization of emerging embryonic cell types based on transcriptional signatures, rather than relying on the definitive morphological and/or functional features of their mature counterparts.

### Cell type equivalencies across developmental and evolutionary time

Such molecular classification schemes echo recent conceptual frameworks that aim to categorize ‘cell types’ across developmental and evolutionary time scales, irrespective of morphology or function [2]. If, however, we consider a ‘cell type’ to be primarily defined by the expression of distinct regulatory programs, then detection of program activities can substantially precede our ability to distinguish morphological or functional specializations. Indeed, our sub-clustering and module analyses across developmental time reveal the appearance of certain prospective cell types long before they become morphologically distinct. For example, already at stage HH25 we recover clear gene expression signatures reminiscent of the future periskeletal nsCT, even though prominent cartilage anlagen have yet to form (Fig. 2d, Fig. 3b). As such, it suggests an early lineage priming, without necessarily implying a definite switch in cell fate or clear morphological distinctions. In agreement with this, our *ZFHX3*-containing module Darkgreen appears to be the most basic and least specific of the co-expression modules that coincide with the nsCT population. We detect its activity at all three time points, marking the prospective nsCT as well as parts of the *PRRX1*-positive mesenchymal progenitor population (Fig. 5c,f). Only later do more mature and restricted nsCT sub-divisions and their corresponding co-expression modules occur, as exemplified by the activity of module Darkgrey and some of its members known to be involved in the formation of periskeletal tissues and tendon attachment sites (Fig. 5a,d) [38, 39].

Moreover, combining such transcriptome-based ‘cell type’ classification schemes with comparative scRNA-seq datasets allows for a molecular assessment of homologous cell types between species, across evolutionary time scales [40, 41]. This has important implications when trying to elucidate the impact of cell type-specifying gene regulatory networks on pattern formation and diversification at its relevant cellular scale. Namely, how progenitor populations exactly perceive and process patterning-relevant cues can be modulated by species-specific alterations in the respective cell type-specifying networks. In this context, it is worth noting that we detect *RSPO3* as one of the main markers of the chicken AER (Fig. 2d, Additional file 2). R-spondins, a family of secreted ligands involved in WNT-signaling, have previously been implicated in AER maintenance and control of limb outgrowth. However, in mammals only *RSPO2*, and not *RSPO3*, seems to be implicated in AER function [42–44]. Similarly, species-specific modifications in the gene regulatory networks driving skeletal cell type maturation have been reported [45, 46]. Together with recent scRNA-seq studies in other vertebrate model organisms [30, 47, 48], our dataset now opens new avenues for a comprehensive assessment of molecular similarities and divergences in patterning-relevant cell populations of the developing limb, across all major tetrapod clades.

### Digit growth and patterning at cellular resolution

Variations in digit number, size and individual digit patterns in the autopod skeletal structure reflect functional specialization of tetrapod hands and feet. During development, condensations of mesenchymal cells first give rise to early skeletogenic progenitors, to then differentiate into distinct skeletal lineages such as chondrocytes, osteocytes or synovial joint cells [49–51]. However, unlike for skeletal elements at more proximal locations of the limb, individual phalanx condensations are sequentially added and expanded at the distal tip of each forming digit, through proliferation of an evolutionary conserved progenitor population [22, 23, 52]. Hence, identifying regulators of growth rates, as well as for the relative temporal sequence at which the different skeletal cell types are specified, becomes paramount when trying to understand digit-specific phalanx patterns [25, 53].

Early autopod outgrowth, and later digit elongation, is controlled through complex signaling interactions at the distal margin of the limb, involving the concerted action of FGFs, BMPs and WNTs [reviewed 5]. Coinciding with this distal domain, we identify a distinct sub-population of mesenchymal cell types in all of our samples, marked by elevated activity of module Magenta with *TFAP2B*, *WNT5A* and high BMP signaling (Fig. 3c-f). Certain module members have been functionally implied in regulating autopod growth and digit elongation [24, 54–56], yet others remain completely unexplored in this context.

Moreover, we identify distinct sub-populations of interdigit mesenchyme cells in our HH29 and HH31 samples, with four associated gene co-expression modules (Fig. 4a-h). Module Olivegreen contains SNAI and ID genes, known to be expressed in interdigits, and likely relates to the various BMP-driven processes in this tissue [57–62]. On the other hand, module Orange is dominated by *RDH10*, implicated in mouse interdigital apoptosis [63]. Before its apoptotic disappearance at later stages of development, interdigit mesenchyme is known to instruct the specific phalanx-formulas of its anteriorly adjacent digit [24, 25]. Moreover, we manage to spatially attribute a distinct co-expression module (Midnightblue) to interdigit 4, i.e. posterior to a digit with known regulatory individualization in tetrapods [64].

Finally, across all developmental time points we sampled, we identify skeletogenic cell populations. At those stages, the forming skeletal elements still consist exclusively of early progenitors, maturing chondrocytes, and developing synovial joints. Accordingly, we only find three distinct sub-populations, associated with two co-expression modules. Module Red shows enrichment for many canonical markers of chondrocyte maturation (Fig. 6e) [45, 51]. On the other hand, genes in module Turquoise do not, for the most part, evoke a classical chondrogenic transcriptional profile (Fig. 6d). Again, this module might rather reflect an early transcriptional priming, only this time towards the skeletogenic lineage. In agreement with this, we only detect low expression levels for the canonical early skeletogenic marker *SOX9* in HH25 sub-cluster 4 (Fig. 2d), which itself is specifically enriched for Turquoise activity. Likewise, our synovial joint-like HH29 sub-cluster 17 shows high activity for Turquoise, while excluding the more mature chondrocyte module Red (Fig. 6g-i).

## Conclusion

Our single-cell transcriptomic atlas provides a comprehensive genomics resource to study chicken limb development in unprecedented detail. Thereby, it complements a classical experimental model of vertebrate pattern formation with molecular data at cellular resolution. We curate molecular catalogues to provide an in-depth description of the embryonic autopod, through the assembly of cell population-specific lists of candidate marker genes. Combined with the power of viral overexpression screens and recent CRISPR/*Cas9* genome modifications technologies, this resource will provide a roadmap for the functional elucidation of cell type specification programs in patterning-relevant populations. Moreover, by constructing cell population-specific gene co-expression modules, we provide a tool to follow tissue dynamics across developmental and evolutionary time scales. Thereby, it will enable insights into the molecular underpinnings of homologous cell types across all major tetrapod clades, and their ensuing developmental impact on pattern formation and diversification in the vertebrate autopod.

## Methods

### Tissue sampling

We collected tissue samples from embryonic hind limbs at different developmental stages (Fig. 1,a-c). Limbs were dissected in cold PBS, and chopped coarsely with a razorblade. Dissociation into single cells was done using 2.5% trypsin in DMEM and incubation for 15 minutes at 37°. Occasional mechanical shearing by careful pipetting was applied during the incubation time.

### scRNA-seq library preparation

Single-cell suspensions of samples HH25 and HH31 were fed into a *10X Genomics Chromium* Single Cell System (*10X Genomics*, Pleasanton, CA, USA) aiming for a concentration of 4000 cells per microliter. Cell capture, cDNA generation, preamplification and library preparation were done using *Chromium Single Cell 3’ v2* Reagent Kit according to the manufacturer instructions. For stage HH29 the cells were processed with the *DropSeq* method according to the original protocol [26]. Once the cDNA was obtained from all the samples, the sequencing proceeded on Illumina *NextSeq 500* platforms as recommended by the developers at 75bp and an average depth of 400 million reads per sample.

### Data processing

Using either the *Cell Ranger* software v2 (*10X Genomics*) or the *DropSeq* pipeline v1 (https://github.com/broadinstitute/Drop-seq/releases) we performed base calling, adaptor trimming, mapping to the chicken ENSEMBL genome assembly and annotation Gallus_gallus-5.0 [65], de-multiplexing of the sequences and generation of the gene / cell count matrices.

Filtering thresholds for mapped data were adapted for each sample, depending on the different library complexities. Cells with an UMI count of more than 4 times the sample mean or less than 20% of the sample median were filtered out, cells with a mitochondrial or ribosomal contribution to UMI count of more than 10% were also filtered out. Using the R package Seurat v2.3.2 [66] the UMI counts were then Log-normalized and any variation due to the library size or mitochondrial UMI counts percentage was then regressed via a variance correction using the function ScaleData.

The cell cycle stage of each cell was inferred using the R package SCRAN [67] and gene pairs that covariate with cell cycle stages in mouse [68]. The gene pairs were translated to orthologous chicken genes [69] and a cell cycle stage score was obtained cell-wise for stages S, G1 and G2/M, the difference between the G2/M and S scores (δG2M/S) was calculated to be accounted for in later steps.

### Dimensionality reduction and visualization

Significant principal components were determined for each sample as those falling outside of a Marchenko-Pastur distribution [35]. A dimensionality reduction step was carried out, using the t-SNE algorithm [28] to visualize the data and clustering of the cells based on transcriptomic similarities. The cells were clustered using the Louvain method for community detection from large networks and the Jaccard similarity coefficient to compare similarity and diversity of the sets, implemented in the FindClusters function in Seurat using data which was additionally variance-corrected for δG2M/S. A first, broad, clustering step was done using a resolution of 0.4 for samples HH31 and HH29 and 0.5 for HH25; a second clustering was done to find sub-clusters within the data, this time using resolutions of 1.4 and 1.1 for the corresponding samples. All clustering steps were done using a k number of 20 and the significant principal components of the sample.

### Differential expression analysis

Differential expression analyses based on the negative binomial distribution were performed with Seurat, using the δG2M/S as a covariate and only genes expressed in at least 15% of any compared population (Additional files 2-4); genes expressed in at least 25% of the cells and showing differences with a log fold-change > 0.5 and an adjusted p value < 0.05 were used for GO analyses. To find expression signatures for every cell cluster, in a first step, a phylogenetic tree was obtained for the cell clusters in each sample; all directly paired clusters were tested for differential expression. Any pair of clusters with less than 15 differentially expressed genes were collapsed recursively. In a second step, specific genes for each cluster were obtained contrasting each cluster against the rest of the cells in their sample. To find genes differentially expressed genes between the interdigit clusters (Fig. 4j), we compared each of the sub-clusters against the rest of the cells in the other two clusters.

Marker genes for digit/interdigit 3 and 4 were defined using the DESeq2 R package v1.20.0 [70]. We analyzed bulk RNA data sets of digit/interdigt 3 and 4 from stage HH28/29 and HH31 of a previous study [34]. After normalization based on size factors and dispersion, we performed the differential expression analysis using a Wald test and the contrast design ∼Stage+Digit to use the different stages as pseudo-replicates of the digit. We filtered for differential expression with a p-value < 0.05. For visualization, we subtracted the fold changes of early and late stages and plotted a heatmap using heatmap3 R package v1.1.1 [71] using hierarchical clustering of the genes.

### Weighted co-expression analyses

A weighted correlation network analysis was done using the WGCNA R package v1.6.6 [32]. Using the function FindVariableGenes from Seurat, we calculated the genes with high variation (dispersion > 0.5) across all the cells in sample HH29, and were subsequently used in WGCNA. Adjacencies and signed topological overlaps were calculated with an inferred soft-thresholding power of 8. A hierarchical tree was constructed using the “average” method and then cut using the “tree” method at height 0.9957 and minimum module size of 15. The eigengenes of the resulting modules, as well as the membership and a Correlation Student p value of the membership of each gene to its module were calculated. All genes not significantly (p value >0.01) correlated with any module were discarded. The process was repeated recursively, until all genes were significantly associated with a module; the only change made in every iteration was the module minimum size, set to the smallest that would yield at least the same number of modules as the first analysis.

The output of WGCNA was exported to the Cytoscape v3.7.0 software [72] where the node size was coded to represent the membership, and the edge thickness and color intensity to represent the weights of each gene-pair coexpression. For visualization purposes, the scales of thickness, color and size were made relative to the minima and maxima found in each network. Furthermore, a transparency gradient was added to the edges, which was scaled to hide unimportant edges and avoid edge saturation, the threshold was always adjusted to make visible at least one edge per node. In only one case (module midnightblue), an edge with an outlier weight was coded to be red and thicker than any other edge, and the color/size re-scaled to the second highest weight.

### Gene Ontology

Gene Ontology analyses were conducted with the R package *limma* [73]. We used the list of genes in the expression signature of each computed cell cluster, and the genes members of each co-expression module as input. For each case we used all the genes detected in the corresponding sample as the contrast universe.

### *In situ* hybridization

Probes for *CRABP-I* and *COL1A2* were described previously [38]. Primers for the *ZFHX3* probe were designed using primer3 [74]. An AA overhang and an *EcoRI* restriction site were added to each of the primers at the 5’ end. ZFHX3 (fw: [5’-<u>AAGAATTCAGC</u>CGTACCGGGTGCAATGAGC-3’], rev: [5’-<u>AAGAATTCAGCG</u>CTTCCTCTTCCCGTAGAGC-3’]). *In situ* hybridization was performed using standard protocols [75]

## Abbreviations

EvoDevo: Evolutionary developmental biology
LPM: Lateral plate mesoderm
AER: Apical ectodermal ridge
HH: Hamburger-Hamilton stages
UMIs: Unique molecular identifiers
tSNE: t-distributed stochastic neighbor embedding
GO: Gene ontology
nsCT: Non-skeletal connective tissue
TFs: Transcription factors
scRNA-seq: Single-cell RNA sequencing

## Declarations

### Availability of data and materials

All data generated or analyzed during this study are included in this published article and its supplementary information files. Raw sequencing data has been deposited in the SRA (accession numbers: TBD)

### Authors’ contributions

PT conceived and designed the study. CF, OP and PT conducted the scRNA-seq experiments. CF conducted data analyses and *in situ* experiments. CF and FS conducted the bulk RNA-seq re-analysis. CF and PT drafted the manuscript. All of the authors read and approved the final manuscript.

### Ethics approval and consent to participate

In accordance with Swiss national guidelines (Swiss Animal Protection Ordinance; TSchV, chapter 6, Art. 112), no formal ethics approval was required, as all experiments were carried out prior to the third trimester of incubation.

### Consent for publication

Not applicable.

## Acknowledgements

Calculations were performed at sciCORE (http://scicore.unibas.ch/) scientific computing center at University of Basel. CF and PT wish to acknowledge Katja Eschbach and Christian Beisel for help with *10X Genomics Chromium* and sequencing. OP and PT thank Tyler Burks for help with *DropSeq* experiments. PT and OP would like to acknowledge the generous support of Cliff Tabin and Aviv Regev, in whose labs this project was initiated (with help of NIH grant HD03443 to Cliff Tabin).

---

**Fig. S1.**
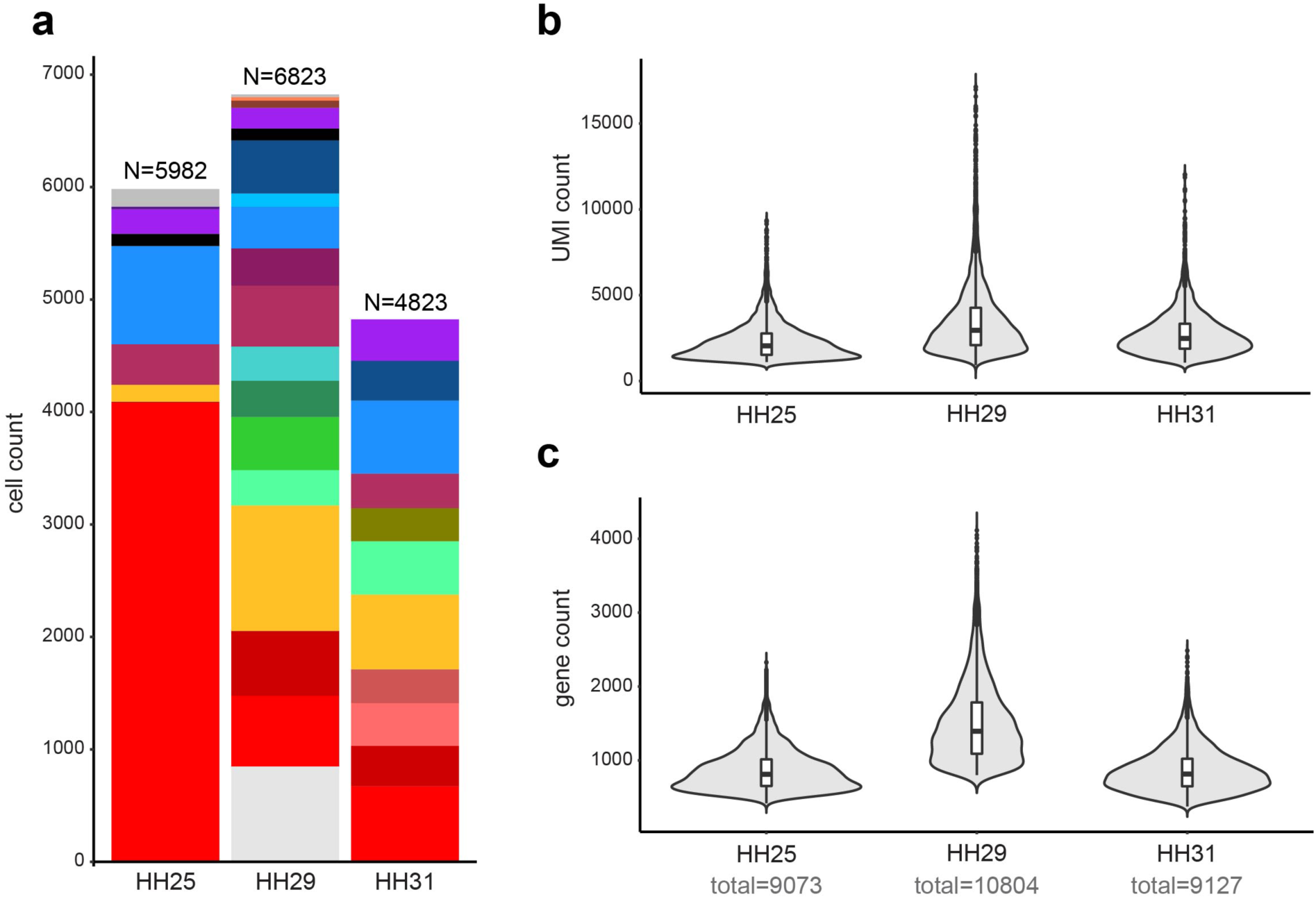
Sample compositions and data statistics. **(a)** Cellular composition of the samples and datasets, color code corresponds to Fig. 2a-c. **(b)** UMI count distributions across the samples. **(c)** Gene count distributions across the samples.

**Fig. S2.**
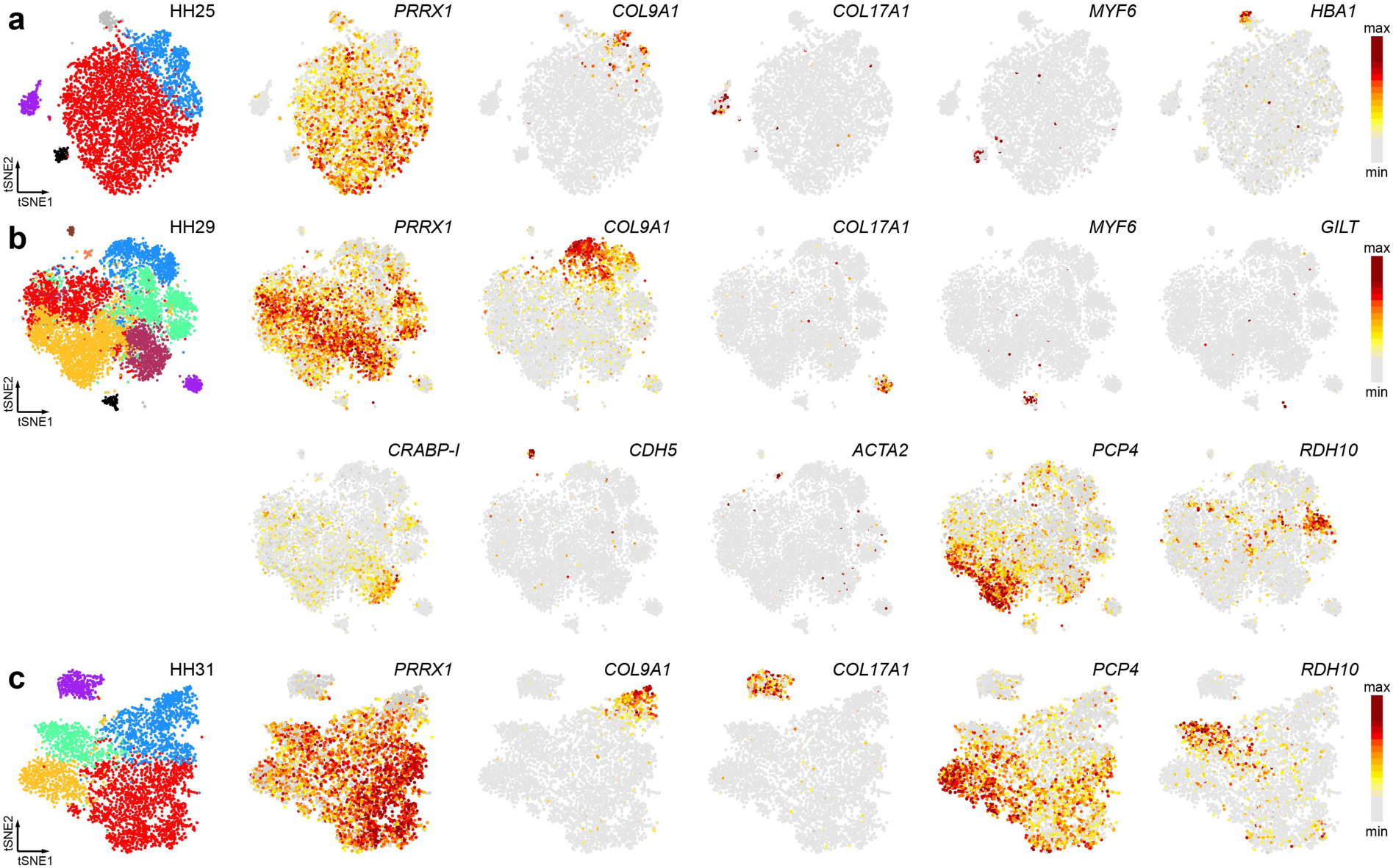
Expression patterns of marker genes. Related to Fig. 1. Normalized expression patterns of selected genes to identify the different cell populations in our broad clustering, plotted on the tSNEs from sample **(a)** HH25, **(b)** HH29 and **(c)** HH31.

**Fig. S3.**
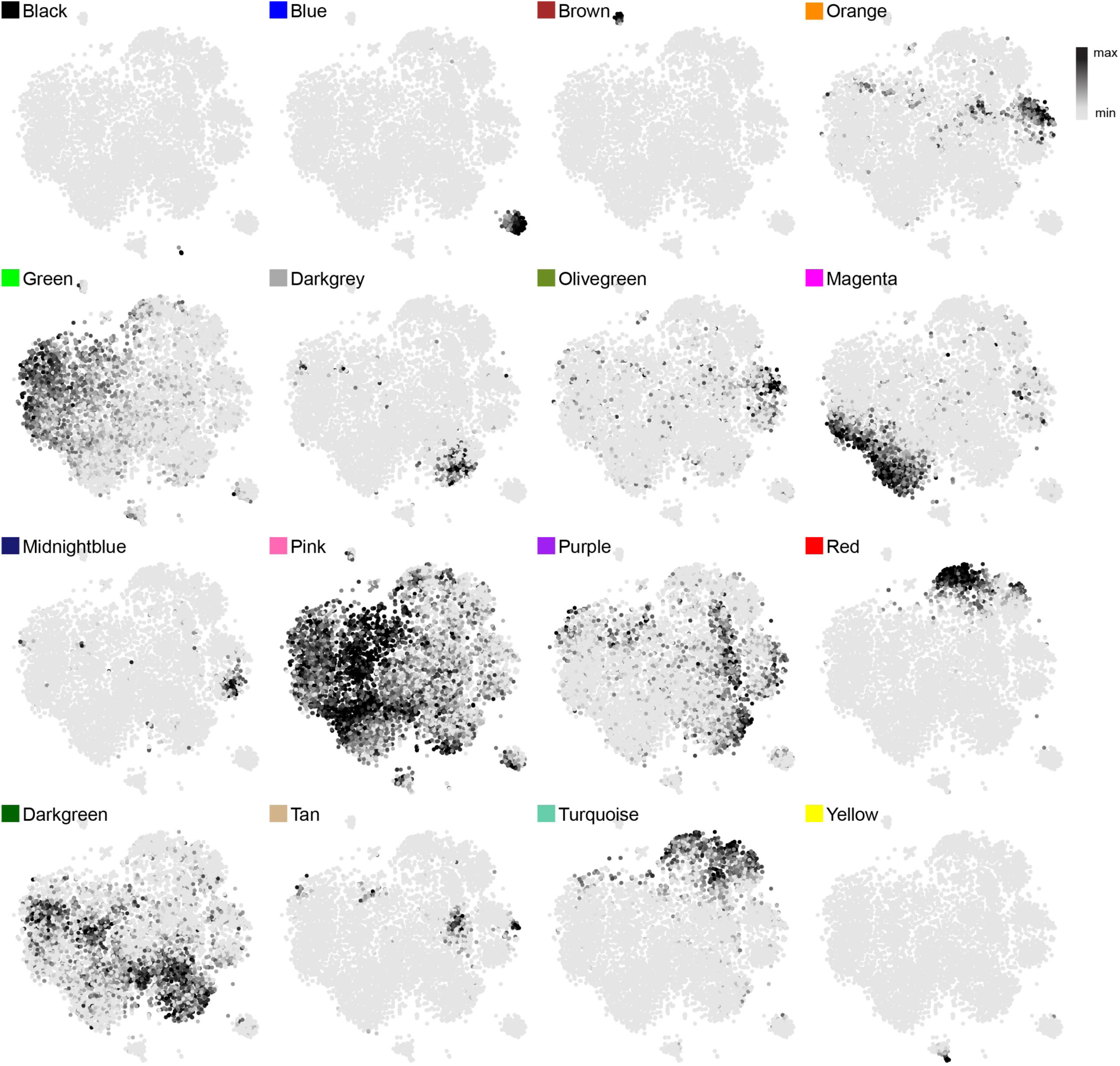
Co-expression modules expression patterns. Related to Fig. 3. Average expression of each WGCNA co-expression module on the tSNE of sample HH29.

## Additional file 1

### XLSX

#### Genes with enriched expression per cell population in sample HH25

Genes enriched in the different cell clusters, calculated to be differentially expressed between each cell cluster and the rest of the cells in the sample. **p_val**: originally calculated p value; **avg_logFC**: average log fold-change relative to the rest of the cells; **pct.x**: percentage of cells in the focus cluster expressing the gene; **pct.rest**: percentage of cells in the rest of the clusters expressing the gene; **p_val_adj**: p value adjusted for multiple testing; cluster: cluster number in the main text and figures; **gene**: ENSEMBL gene identifier; **name**: gene symbol, or name when available; **enrichment**: ratio of pct.x: pct.rest.

## Additional file 5

### XLSX

#### Co-expression modules and their genes

Genes part of the different co-expression modules. **nodeName**: ENSMBL identifier of the genes part of the module; **altName**: gene symbol, or name when available; **membership**: membership to the module.

